# High-frequency oscillations in the internal globus pallidus: a pathophysiological biomarker in Parkinson's disease?

**DOI:** 10.1101/2020.06.16.144477

**Authors:** Luke A Johnson, Joshua E Aman, Ying Yu, David Escobar Sanabria, Jing Wang, Meghan Hill, Rajiv Dharnipragada, Remi Patriat, Mark Fiecas, Laura Li, Lauren E Schrock, Scott E Cooper, Matthew D Johnson, Michael C Park, Noam Harel, Jerrold L Vitek

## Abstract

Abnormal oscillatory neural activity in the basal ganglia is thought to play a pathophysiological role in Parkinson’s disease. Many patient studies have focused on beta frequency band (13-35 Hz) local field potential activity in the subthalamic nucleus, however increasing evidence points to alterations in neural oscillations in high frequency ranges (>100 Hz) having pathophysiological relevance. Prior studies have found that power in subthalamic high frequency oscillations (HFOs) is positively correlated with dopamine tone and increased during voluntary movements, implicating these brain rhythms in normal basal ganglia function. Contrary to this idea, in the current study we present a combination of clinical and preclinical data that support the hypothesis that HFOs in the internal globus pallidus (GPi) are a pathophysiological feature of Parkinson’s disease. Spontaneous and movement-related pallidal field potentials were recorded from deep brain stimulation (DBS) leads targeting the GPi in five externalized Parkinson’s disease patients, on and off dopaminergic medication. We identified a prominent oscillatory peak centered at 200-300 Hz in the off-medication rest recordings in all patients. High frequency power increased during movement, and the magnitude of modulation was negatively correlated with bradykinesia. Moreover, high frequency oscillations were significantly attenuated in the on-medication condition, suggesting they are a feature of the parkinsonian condition. To further confirm that GPi high frequency oscillations are characteristic of dopamine depletion, we also collected field potentials from DBS leads chronically implanted in three rhesus monkeys before and after the induction of parkinsonism with the neurotoxin 1-methyl-4-phenyl-1,2,3,6 tetrahydropyridine (MPTP). High frequency oscillations and their modulation during movement were not prominent in the normal condition but emerged in the parkinsonian condition in the monkey model. These data provide the first evidence demonstrating that exaggerated, movement-modulated high frequency oscillations in the internal globus pallidus are a pathophysiological feature of Parkinson’s disease, and motivate additional investigations into the functional roles of high frequency neural oscillations across the basal ganglia-thalamocortical motor circuit and their relationship to motor control in normal and diseased states. These findings also provide rationale for further exploration of these signals for electrophysiological biomarker-based device programming and stimulation strategies in patients receiving deep brain stimulation therapy.

## Introduction

Numerous studies suggest that abnormal neural oscillatory activity in the basal ganglia plays a pathophysiological role in Parkinson’s disease (PD). For example, recordings of local field potentials (LFPs) in the subthalamic nucleus (STN) of patients undergoing deep brain stimulation (DBS) have shown relationships between beta frequency band (13-35Hz) oscillatory features to parkinsonian motor signs such as akinesia, bradykinesia and rigidity (Brown, 2003; Kühn *et al.*, 2006, 2009; Ray *et al.*, 2008; Shreve *et al.*, 2017). Increasing evidence, however, suggests that neural oscillations in frequency ranges outside the beta band also have pathophysiological relevance in PD (Petersson *et al.*, 2019). In the STN alterations in high frequency oscillations (HFOs, >100Hz) have been purported to reflect changes in the motor state, with studies finding HFO band power increases during movement and dopaminergic treatment causes HFO peaks to emerge (Foffani *et al.*, 2003) and/or shift towards faster frequencies (e.g. from 200-300Hz to 300-400Hz) (López-Azcárate *et al.*, 2010; Özkurt *et al.*, 2011). As such, it is suggested that HFO activity in the STN has a pro-kinetic role and may be important for normal motor control (Foffani *et al.*, 2003).

Despite the fact that the internal globus pallidus (GPi) is increasingly being targeted for DBS in PD patients, there are relatively few reports exploring LFP activity in the pallidum (Priori *et al.*, 2002; Silberstein *et al.*, 2003; Tsiokos *et al.*, 2013, 2017; AuYong *et al.*, 2018; Aman *et al.*, 2020). To date only one group has published PD patient studies that focused on GPi HFOs (Tsiokos *et al.*, 2013; AuYong *et al.*, 2018), finding movement-enhanced peaks in oscillatory activity centered around 230Hz based on intraoperative GPi DBS lead recordings. These studies raise questions about the pathophysiological relevance of GPi HFOs and whether they are a prokinetic signal important for normal basal ganglia motor function (Tsiokos *et al.*, 2013). To address these questions, we collected spontaneous and movement-related LFPs from DBS leads in GPi from externalized PD patients on and off dopaminergic medication, as well as from rhesus monkeys before and after the induction of parkinsonism with the neurotoxin 1-methyl-4-phenyl-1,2,3,6 tetrahydropyridine (MPTP). We hypothesized that GPi HFO activity would have similar prokinetic characteristics as observed in the STN of PD patients, with HFO amplitude positively correlated with dopaminergic tone and enhanced during voluntary movements (Foffani *et al.*, 2003; López-Azcárate *et al.*, 2010). Contrary to this hypothesis, the present study provides compelling evidence that rather than a feature important for normal motor control, exaggerated, movement-modulated HFO activity in the GPi is in fact a pathophysiological feature of the parkinsonian condition. These findings improve our understanding of the pathophysiological role of basal ganglia high frequency oscillatory activity in Parkinson’s disease and support further exploration into their functional role in the development of specific motor signs, as well as how these signals may inform DBS device programming (Horn *et al.*, 2017; Tinkhauser *et al.*, 2018) or the design of electrophysiological biomarker-based adaptive stimulation approaches (Meidahl *et al.*, 2017) in patients receiving GPi DBS.

## Materials and methods

### Data collection in PD patients

All patient procedures were approved by the University of Minnesota Institutional Review Board (#1701M04144) with consent obtained according to the Declaration of Helsinki. Five patients (two female, three male) with idiopathic Parkinson’s disease approved for GPi DBS by our consensus committee were consented for externalization and enrolled in the study. All patients were implanted unilaterally. Demographics and total pre-surgery UPDRS-III (motor) scores for each patient are provided in Table 1.

**Table 1:** Patient Demographics and Clinical Ratings of PD Motor Signs Off/On Levodopa

| Patient | Age/Sex | Disease<br>duration (yrs) | DBS<br>side | UPDRS-III<br>OFF | UPDRS-III<br>ON |
| --- | --- | --- | --- | --- | --- |
| 1 | 60 / F | 10 | Right | 39 | 17 |
| 2 | 52 / M | 6 | Right | 39 | 16 |
| 3 | 55 / M | 6 | Right | 17 | 12 |
| 4 | 72 / M | 10 | Left | 56 | 29 |
| 5 | 63 / F | 7 | Left | 44 | 19 |

Details of the surgical procedures for GPi DBS implantation and lead externalization are described in detail in a previous publication (Aman *et al.*, 2020). Briefly, patients underwent standard 3T MRI (all patients) and a high resolution 7T MRI (4/5 patients, excluding Pt 3) for direct targeting and postoperative lead localization (Duchin *et al.*, 2018; Patriat *et al.*, 2018). Intraoperative electrophysiological mapping techniques (Vitek *et al.*, 1998) were used to identify the sensorimotor region of GPi for implantation. In all patients a directional “1-3-3-1” electrode was used (4/5 patients: Abbott Infinity model 6172, illustrated in **Figure 1C**; 1/5 patients: Boston Scientific Vercise Cartesia model DB-2202-45; both leads had 1.5 mm contact height with 0.5 mm vertical spacing). After implantation the lead extension was tunneled to a subcutaneous pocket in the chest and then connected to another extension externalized at the abdomen (Aman *et al.*, 2020). Externalized components were secured and protected with a water-proof barrier dressing and patients were discharged to home to recover. Externalization recordings occurred 4-8 days later, allowing some time for reduction of microlesion effects that can occur following lead placement (Koop *et al.*, 2006; Vitek *et al.*, 2020).

**Figure 1.**
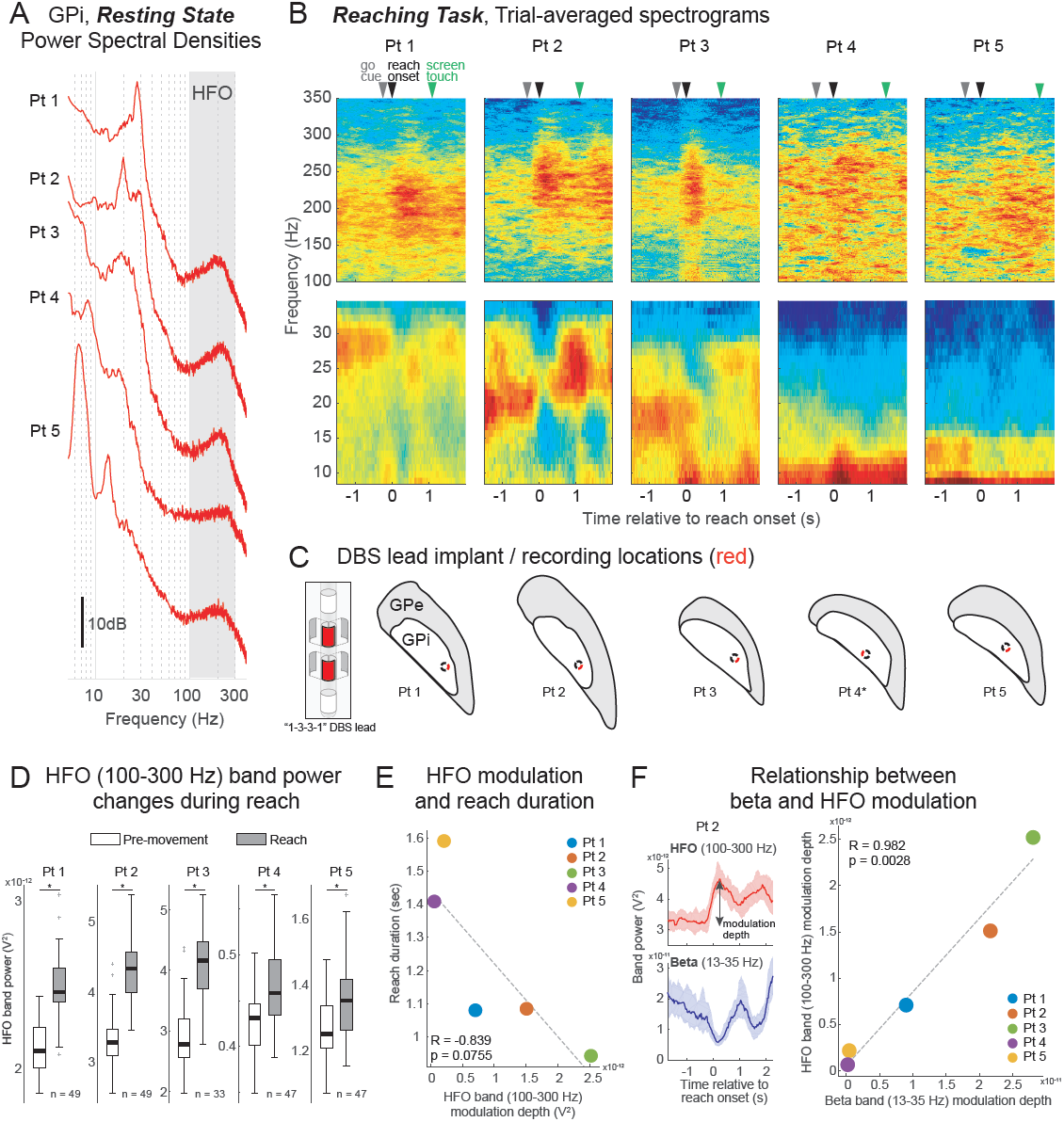
Resting state and movement-related high frequency oscillations recorded from DBS leads in the GPi of PD patients. **(A)** Oscillatory activity observed in the GPi of five externalized DBS patients, derived from rest recordings. Median power spectral density shown (see Methods). **(B)** GPi oscillatory activity during a touchscreen reaching task. Trial-averaged spectrograms aligned to reach onset (t = 0) in higher (100-350 Hz, *upper panels*) and lower (8-35 Hz, *lower panels*) frequency ranges for each patient are shown. Median times of go cue and initial target touch on the touchscreen relative to reach onset are indicated by gray, green and black arrows, respectively. Numbers of reach trials are shown in panel D. **(C)** *Left panel:* schematic of an Abbott directional “1-3-3-1” lead (left panel), illustrating in red that bipolar paired recordings from vertically adjacent segments were used in this study. *Right panels:* DBS lead implant locations for each patient, estimated from preoperative MRI and postoperative CT scans. Axial reconstructions are shown, with recording segment direction indicated in red. Recordings were made in left GPi in Pt 1, Pt 2 and Pt 4 and in right GPi in Pt 3 and Pt 5; right GPi images were mirrored for visualization. *Posterolateral segments were chosen for analysis in all patients except Pt 4 (see Methods). **(D)** HFO (100-300 Hz) band power calculated in a pre-movement period 1 sec duration prior to go-cue compared to band power calculated in a 1 sec duration period beginning with reach onset. In every patient there was a significant increase in HFO band power during reach (Wilcoxon Rank-sum (WRS) test, * p<0.05). **(E)** Relationship between HFO band power modulation depth during reach and reach duration. Across patients there was a strong negative linear relationship between HFO band power modulation and reach duration which did not reach statistical significance (Pearson, Rho = -0.839, p = 0.0755). **(F)** Relationship between beta (13-35 Hz) and HFO band power modulation during reach. *Left panel:* Band power over time, relative to reach onset, from Pt 2, illustrating concomitant modulations in beta and HFO band power. *Right panel:* Across patients, there was a significant linear relationship between the magnitude of beta power decrease and HFO power increase (Pearson, Rho = 0.982, p = 0.0028).

Spontaneous and task-related LFP activity were recorded from the DBS lead over the course of two days while residing in the University of Minnesota Health Clinical Research Unit. The externalized lead was connected to an ATLAS Neurophysiological System (NeuraLynx, Inc) via an adapter cable. Neural data were acquired (EEG scalp contacts used for reference and ground) and digitized at 24kHz for offline analysis. Movement kinematics during the reach task were collected simultaneously with lead LFPs using Delsys Trigno Legacy inertial measurement unit (IMU) wireless sensor placed on the contralateral hand (Delsys, Inc.).

Spontaneous resting state data were collected for five minutes while the patient remained seated in their hospital bed with instructions to remain still and look at a predetermined fixed point on the wall directly in front of them. Movement data were collected while patients performed a touchscreen reaching task using the arm contralateral to the implanted DBS lead. Trials began with the hand on a digitized home button located 45 cm from a touchscreen monitor. After a randomized variable 3-4 sec delay following the start of a trial, a 1.27 cm hollow circle (target) appeared on the center of the touchscreen along with a 5 cm square box directly to the left of the circle (10 cm). The appearance of the circle and square was the patient’s “go cue”. Subjects were instructed to touch and drag the circle into the square box as quickly and accurately as possible and then return to the home button (50 total trials). Initiating a movement prior to the go cue, or failing to complete the task within a 10 s window resulted in the trial being removed from further offline analysis.

Off-medication data were collected after overnight withdrawal of dopaminergic medication. On medication (L-dopa) data were collected approximately 60 minutes after administration of oral medication and after the patient confirmed their on-medication status. At the conclusion of the study patients returned to the hospital for removal of the percutaneous extension and placement of the IPG. A movement disorders clinician performed DBS programming approximately 4-6 weeks after IPG placement per standard clinical care.

DBS lead locations in the GPi were estimated based on information obtained during intraoperative electrophysiological mapping as well as co-registered preoperative MRI and postoperative CT scans (see (Duchin et al., 2018; Patriat *et al.*, 2018) for details). The orientation of the DBS lead and relative direction of individual segments for each patient were derived from the fiducial marker on the lead, in combination with the unique artifact characteristics of the segments, using a modified version of the DiODe algorithm (Hellerbach *et al.*, 2018) combined with information extracted from fluoroscopy and X-ray images acquired intraoperatively.

### Data collection in monkeys

All monkey procedures were approved by the University of Minnesota Institutional Animal Care and Use Committee and complied with United States Public Health Service policy on the humane care and use of laboratory animals. Three adult female rhesus macaques (*Macaca mulatta*, K (13 years), J (16 years) and P (18 years)) were chronically implanted with scaled-down versions of human DBS leads (0.5 mm height ring contacts, 0.5 mm inter-contact spacing, 0.625 mm diameter, NuMED, Inc.) targeting sensorimotor GPi, based on microelectrode mapping procedures analogous to what is done clinically (Vitek *et al.*, 1998). These methods are described in detail in our previous publications (Hashimoto *et al.*, 2003; Elder *et al.*, 2005). Animals were instrumented with additional electrophysiology hardware targeting other brain areas but these were not used in the present study.

Neural data were acquired using a TDT neurophysiology workstation (Tucker Davis Technologies) and digitized at ≈24kHz for offline analysis. Spontaneous awake resting state data were collected while the animal was seated in a primate chair for ≈5 min per recording session as described in our previous publication (Escobar Sanabria *et al.*, 2017). Once data were collected in the naïve state, animals were rendered parkinsonian with a series of intramuscular injections (0.3-0.8 mg/kg per injection) of the neurotoxin 1-methyl-4-phenyl-1,2,3,6 tetrahydropyridine (MPTP) that produces a model system which closely mirrors the human PD condition (Burns *et al.*, 1983; Vitek and Johnson, 2019). In Monkey J, additional data were collected after a subsequent intracarotid injection (0.4 mg/kg). In this monkey we also report data collected during a reaching task in normal, mild (after intramuscular injections) and moderate (after intracarotid injection) PD conditions. A trial began with the hand positioned on a start pad, after which a food reward was presented directly in front of the monkey. The trial was completed after the monkey reached to the reward and returned it to its mouth. Movement kinematics during the reach task were collected simultaneously with lead LFPs using a motion capture system (Motion Analysis Corp.) that tracked position of a reflective marker placed on the monkey’s wrist.

### Data analysis and statistics

All analyses were performed using customized scripts in MATLAB (MathWorks, 2016). Raw signals were digitally bandpass filtered (0.5 - 500 Hz) and down sampled (≈3 kHz). LFP activity was extracted via bipolar montage (i.e. signal subtraction) of vertically adjacent DBS contacts within the GPi. In PD patients, only the segmented contacts of the 1-3-3-1 directional lead were used; the bipolar pair that faced the posterolateral “sensorimotor” territory was chosen for subsequent analysis (see **Figure 1C**, Aman *et al.*, 2020). In patient 4 the anteromedial bipolar pair was used for analysis since this pair had higher signal to noise ratio than the other contact pairs.

Resting state data were divided into 15 sec epochs and power spectral densities (PSDs) for each epoch were calculated using Welch’s method, with 2^14^ points (frequency resolution ≈0.2 Hz) in the discrete Fourier transform, a 2^14^ point Hamming window and an overlap of 50%. Median PSD and 25^th^-75^th^ percentiles across all epochs are shown in the figures. To compare high frequency oscillatory (HFO) activity between off/on medication conditions (patients) and between naïve/MPTP conditions (monkeys), PSD values for each 15 sec epoch were summed in the 100-300Hz range to compute the total power in this band. The distributions of power in each condition were compared using the Wilcoxon rank-sum (WRS test, p < 0.05). We recognize that the frequency range of HFOs is not universally defined; we have chosen to define HFOs in the 100-300Hz range empirically based on our data set to encompass the spectral peaks observed in our recordings, typically centered near 200Hz. Off medication data in patients 1, 2, and 5 were included in a previous publication (Aman *et al.*, 2020), focusing on spatial topography of resting and movement-related low frequency (5-40Hz) activity rather than HFOs as in the present study. A portion of resting state data from monkeys J and K (normal condition) and monkey K (parkinsonian condition) was included in a previous publication (Escobar *et al.*, 2017); here we expand the number of datasets and animals, focusing analysis on oscillatory activity in the 100-300Hz range. The total number of rest recording days and 15 sec LFP segments for each animal and across disease states are as follows using format Monkey_condition_(days/segments): K_normal_(4/23), K_PD_(15/140); J_normal_(13/116), J_PD_(21/277); P_normal_(14/96), P_PD_(23/459). In Monkey P, DC shifts in the power spectrums were observed during the experimental period, therefore a Z-score normalization procedure of each LFP time segment was employed prior to spectral analysis (Lofredi et al., 2019).

Reach task data were analyzed using the following methods: For PD patient recordings, the time of reach onset was identified based on the Delsys gyroscope signals (vector sum of x,y,z angular velocities exceeding a threshold of 10°/s after cue presentation). Reach duration was defined as time between reach onset and touch of the target on the touchscreen. LFP data were further down sampled to ≈1kHz and perievent spectrograms aligned to reach onset were computed via the multi-taper method and Chronux toolbox (Bokil *et al.*, 2010) with a frequency resolution of ≈1 Hz and time resolution of 25 ms. Median spectrograms across trials are shown in the figures. To characterize changes in oscillatory activity during reach behavior total power in the frequency ranges 13-35 Hz and 100-300 Hz were calculated by summing perievent spectrograms across frequencies at each time point, and are referred to as beta and HFO band power, respectively. To quantify reach-related changes in band power, mean band power in a pre-movement period 1 sec duration prior to go-cue and in a movement period 1 sec duration beginning with reach onset were calculated for each trial. Distributions between pre-movement and reach were compared using the WRS-test (p < 0.05). Beta and HFO modulation depths were calculated as the maximum minus the minimum of band power during the reach trial. The relationships between reach duration and HFO modulation, and between beta and HFO modulation, were examined by plotting the median values for each patient and performing a Pearson linear correlation analysis. For comparison of reach data in L-dopa and off medication conditions, analysis of high frequency activity was further refined to a frequency range (±75Hz) centered at the peak frequency identified for each patient based on their off-medication PSD. L-dopa condition reach data are presented from 3 patients; in the other two patients L-dopa reach data were not obtained (Pt 2) or were corrupted by artifacts (Pt 3). In patients 1 and 3 there were reach trials that included high amplitude wide band power increases, possibly due to excessive movement artifacts, which were removed from subsequent analysis. The number of reach trials (out of 50 total trials) included in analysis are reported in the figure legends. For monkey task recordings the spectral analysis approach was identical to that just described but with reach onsets identified based on wrist velocity data extracted from a motion capture system (vector sum of x,y,z velocity exceeding a threshold of 30 mm/s after food presentation).

## Data availability

Raw data were generated at the University of Minnesota. Derived data supporting the findings of this study are available from the corresponding author upon reasonable request.

## Results

### High frequency (100-300Hz) oscillations are present in PD patients and increase during movement

We first sought to identify whether high frequency oscillations (HFOs, 100-300 Hz) were present in our PD patient population. Power spectral density (PSD) plots calculated from resting state LFPs illustrate that all five PD patients in this study had prominent peaks in oscillatory activity in the 100-300 Hz range in GPi (**Figure 1A**), centered at 214 ± 22 Hz (mean ± SD). Furthermore, high frequency activity was dynamically modulated during the reaching task (**Figure 1B, *top panel***; median go-cue and screen touch times relative to reach onset indicated by gray and green arrows, respectively). The estimated recording locations within the GPi for each patient are indicated in red in **Figure 1C**.

Significant increases in power in the 100-300 Hz band were present during reach compared to the premovement period (**Figure 1D**), though the magnitude of this increase varied by patient. Patients with greater HFO modulation generally had shorter reach times (**Figure 1B, *top panel***), suggesting that HFOs modulated by movement may have distinct behavioral relevance. The negative correlation between HFO modulation depth and reach duration, although strong across the five patients (**Figure 1E**, Pearson’s linear correlation, Rho = -0.839), did not reach statistical significance (p = 0.0755). There was, however, a significant correlation between movement related changes in HFO modulation and UPDRS-III clinical rating measures of bradykinesia (**Figure 2**, Pearson’s linear correlation, Rho = -0.981, p = 0.0032).

**Figure 2.**
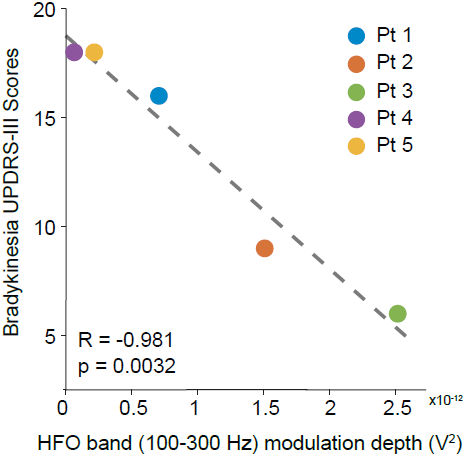
Relationship between movement related changes in HFO power and clinical ratings of bradykinesia. Across patients there was a strong negative linear relationship between HFO band power modulation and UPDRS-III bradykinesia subscores (Pearson, Rho = -0.981, p = 0.0032).

Consistent with numerous studies showing movement related desynchronization of beta band (13-35 Hz) oscillatory activity across brain regions (Engel and Fries, 2010), we found decreased GPi beta activity during reach (**Figure 1B, *lower panel***), though the magnitude of this change also varied by patient. We observed a reciprocal change in magnitude of beta power desynchronization and HFO band synchronization (**Figure 1F, *left panel***), which raised the question of whether there was some relationship across patients between beta and HFO band activity. Indeed, we found a significant linear correlation across patients between the magnitude of the decrease in beta band power and increase in HFO band power that occurred during the movement task (Pearson’s linear correlation, Rho = 0.982, p = 0.0028), i.e. patients with greater HFO synchronization also had greater reach-related beta desynchronization.

### Exaggerated high frequency activity is a pathophysiological feature of parkinsonism

In the previous section we demonstrated that peaks in HFO activity are present in the GPi of PD patients and that movement modulates this activity, but these findings alone are insufficient to address whether HFO activity in the GPi is a pathophysiological feature of the parkinsonian condition. To better address this question, we compared resting state GPi LFPs collected in the off-medication and on-medication (L-dopa) conditions in each patient, as shown in the PSD plots in **Figure 3A**. In every patient dopaminergic medication caused a significant decrease in power in the 100-300 Hz band (WRS test, p < 0.05), with the prominent peak of activity in this range completely abolished in 2/5 patients (Pt 1, Pt 2).

**Figure 3.**
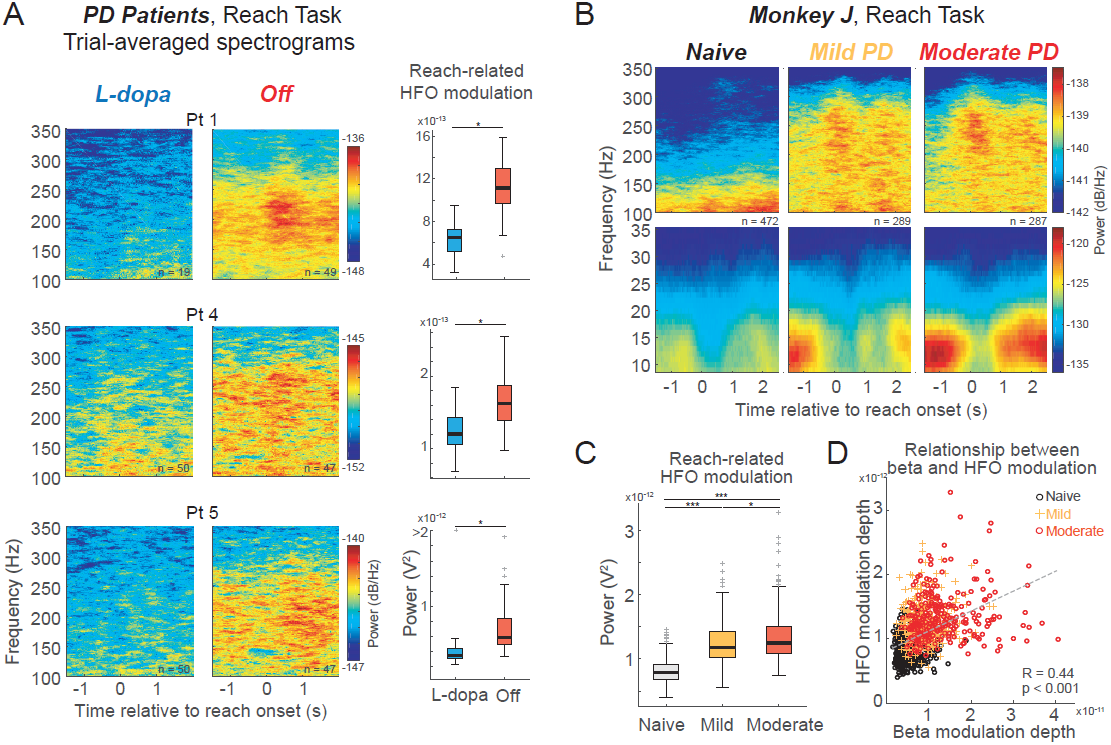
Exaggerated HFO activity is a feature of the parkinsonian condition. **(A)** High frequency oscillations observed in the GPi of five externalized DBS patients were reduced after dopaminergic medication (L-dopa) was administered. Data shown are median power spectral densities from rest recordings in medication off (black) and L-dopa conditions (blue); shaded regions indicate 25-75^th^ percentiles. **(B)** Power spectral densities based on resting state data collected in three monkeys before (naïve, black) and after systemic intramuscular administration of the neurotoxin MPTP (parkinsonian, red), illustrating an emergence of high frequency oscillations in the parkinsonian condition.

To further test the hypothesis that HFO activity is exaggerated in PD, in our preclinical monkey studies we analyzed LFP recordings from chronically implanted DBS electrodes in order to make within-subject comparisons of GPi oscillatory activity before and after induction of parkinsonism using the neurotoxin MPTP. We found distinct peaks in the 100-300 Hz frequency range were not evident in resting state data collected in the naïve condition, but rather emerged in the parkinsonian condition (**Figure 3B**), centered at 238 ± 17 Hz (mean ± SD). The increase in HFO band power was significant in all three animals (WRS test, p < 0.05). Together, patient and monkey data suggest that elevated HFO activity in the GPi is a pathophysiological feature of the parkinsonian condition.

### Movement-related modulation of high frequency activity is characteristic of the parkinsonian condition

The movement related increase in HFO activity presented in **Figure 1B,D** is suggestive of its role in facilitating movement. Yet we found that movement related increases in HFO activity were primarily characteristic of the parkinsonian condition (**Figure 4**). In PD patients, trial averaged spectrograms aligned to reach onset in L-dopa and off conditions illustrate salient synchronization in the 100-300 Hz range primarily in the off-medication condition (**Figure 4A, *left panels***). This is further quantified by reach related modulations in HFO band power, centered on each patient’s HFO peak, which were significantly greater in the off-medication condition (**Figure 4A, *right panels***, WRS-test, p < 0.05).

**Figure 4.** Reach-related modulation of HFO activity occurs predominantly in the parkinsonian condition. **(A)** *Left panels:* GPi oscillatory activity in PD patients collected during the touchscreen reaching task in both medication on (L-dopa) and medication off conditions. Trial-averaged spectrograms aligned to reach onset (time = 0) for each patient are shown. *Right panels:* HFO band power modulation during the reach task in medication on and off conditions. * p<0.05, WRS-test. **(B)** Trial averaged spectrograms from monkey J, in naive, mild (after 3 systemic intramuscular MPTP injections) and moderate (after subsequent intracarotid MPTP injection) parkinsonian conditions. *n* reflects number of reaches in each PD condition. **(C)** Progressive increase in reach-related HFO modulation was observed with increasing parkinsonian severity. * p<0.05, *** p<0.001. **(D)** Significant linear correlation between magnitude of beta modulation depth (desynchronization) and HFO modulation (synchronization) depth across reach trials in all three conditions (Pearson Rho = 0.44, p < 0.001).

LFP data collected during a reaching task in Monkey J before and after MPTP administration provide additional evidence that reach-related synchronization of HFO activity occurs primarily in the dopamine depleted condition (**Figure 4B, *upper panels***). Reach-related HFO modulation depths significantly increased after induction of parkinsonism, and increased further after additional MPTP administration (**Figure 4C**, WRS-test, Bonferroni corrected for multiple comparisons, p < 0.05). Together these patient and monkey data suggest that strong synchronization of HFO activity during movement is not a characteristic of normal basal ganglia function as previously hypothesized (Foffani *et al.*, 2003; Tsiokos *et al.*, 2013) but rather appears to be a feature of neural activity in the GPi of the parkinsonian brain. Consistent with our patient data showing a relationship between beta and HFO band power modulation, there was also a significant positive correlation between these features across conditions within monkey J (**Figure 4D**, Pearson’s linear correlation, Rho = 0.44, p < 0.001)

## Discussion

These data provide the first evidence indicating that exaggerated, movement-modulated high frequency oscillations in the internal globus pallidus are a pathophysiological feature of Parkinson’s disease. This finding is supported by our monkey studies characterizing GPi oscillatory activity before and after MPTP induction of parkinsonism, providing a unique within-subject comparison of naïve and disease conditions not feasible in human studies. Importantly, these monkey data demonstrate that the attenuation of HFO activity in the GPi by levodopa in patients is not a phenomenon unique to medication; rather exaggerated high frequency oscillatory activity in the GPi appears to be characteristic of the parkinsonian, dopamine depleted condition.

### Comparison to previous studies of HFOs in the human basal ganglia

The first report of high frequency oscillations in the basal ganglia of PD patients was by Foffani et al 2003, who described distinct peaks in power spectra centered around 300 Hz in STN LFPs recorded from externalized DBS leads (Foffani *et al.*, 2003). They found a prominent 300 Hz rhythm in only 3/11 recorded nuclei absent dopaminergic medication; after levodopa there was a robust emergence or increase in HFO power in the STN for all subjects. In other studies, HFOs have been more reliably obtained in the STN in PD patients off medication, with both the amplitude and peak frequency strongly modulated by dopamine (López-Azcárate *et al.*, 2010; Özkurt *et al.*, 2011; Hirschmann *et al.*, 2016). These findings are in stark contrast to the robust attenuation of HFO activity we observe in the GPi after levodopa.

Indeed, one of the primary differences between the STN and GPi HFOs appear to be their response to levodopa. In the STN there is marked increase in baseline high frequency power and movement-related modulation of high-frequency power in the on state (Foffani *et al.*, 2003; López-Azcárate *et al.*, 2010), and in some studies there are shifts towards higher HFO peak frequencies (López-Azcárate *et al.*, 2010; Özkurt *et al.*, 2011; Hirschmann *et al.*, 2016). In GPi the reverse is observed, in the sense that significant movement-related modulation of GPi HFO activity is observed primarily in the off-medication condition, and HFO activity is dramatically attenuated after L-dopa administration.

### Relevance of HFO modulation to PD motor signs

Lopez Azcarate et al. 2010 found a correlation between HFO power modulation in the STN and composite bradykinesia & rigidity UPDRS scores, with patients that exhibited greater HFO power increase during movement having less motor impairment (López-Azcárate *et al.*, 2010). They suggest that *“the parameter most closely related to bradykinesia and rigidity is the impairment of movement-related amplitude modulation of the HFOs.”* Interestingly we found a similar relationship between HFO modulation in the GPi and motor impairment in the medication off condition: patients with the greatest reach-related HFO modulation tended to have faster reach times (**Figure 1E)** and there was a significant negative correlation between HFO modulation and UPDRS-III bradykinesia subscores (**Figure 2**). Despite aforementioned differences in the levodopa response of HFOs in STN and GPi, together these findings are suggestive of a common functional role of HFO oscillations in the basal ganglia in the dopamine depleted condition, with dynamic modulation of these signals being important for motor control in the parkinsonian brain. Unlike the STN, however, where modulation of this activity increases with medication, our data from patients on L-dopa and naïve monkeys suggest that, at least in the GPi, these high frequency signals are not characteristic of normal basal ganglia function. Although they may be a prokinetic signal important for motor control of the parkinsonian condition, their prominent presence in the GPi primarily in the dopamine depleted condition may suggest rather that they are reflective of some kind of compensatory process and not a signaling process characteristic of normal brain function.

### Mechanisms underlying GPi HFOs

The cellular mechanisms underlying high frequency oscillations in the basal ganglia are unclear but may reflect synchronous local neuronal firing (Petersson *et al.*, 2019). One hypothesis is that HFOs reflect coordinated bursting across populations of GPi neurons analogous to the coherent neural activity thought to underlie high frequency oscillations in the hippocampus (Buzsaki *et al.*, 1992). Mean discharge rates and bursting characteristics of GPi neurons have been reported to increase in the parkinsonian state (Filion and Tremblay, 1991; Boraud *et al.*, 1998). It is notable that in the study by Filion and Tremblay there was a distinct peak in the mean population interspike interval (ISI) distribution at 4 ms in neurons recorded in the GPi of parkinsonian monkeys, corresponding to an instantaneous firing rate of 250Hz. This is near the 238 ± 17Hz and 214 ± 22Hz mean frequency HFO peaks we observed in our monkeys and PD patients, respectively. Neurons in GPi do not have sustained firing rates above 200 Hz (mean firing was 95 Hz in the study by Filion and Tremblay), but the ISI peak at 4 ms likely reflects the burst firing that is more prominent in the parkinsonian condition, whereas the ISI distribution in normal animals was more diffuse. Firing rates and bursting in the STN are also increased in the parkinsonian condition, though observed mean firing rates are typically lower in the STN than GPi (Bergman *et al.*, 1994; Levy *et al.*, 2001; Du *et al.*, 2018). Nevertheless, increased burst firing and discharge rates are characteristic features of the parkinsonian condition in both the STN and GPi; the potential relationship between these firing patterns and high frequency oscillatory activity observed in LFP recordings in the STN compared to GPi requires further study using high density electrode arrays that can capture LFPs simultaneous with population neuronal spiking activity (Jun *et al.*, 2017). A better understanding of the neuronal mechanisms underlying these pathological oscillations will aid in the design of stimulation strategies optimized to disrupt them or shift them towards a more normal state (Froemke and Dan, 2002; Tass, 2003; Hammond *et al.*, 2007)

### Concluding Remarks

Our data provide new insight into the role of abnormal neural oscillatory activity in the pathophysiology of Parkinson’s disease. These results should motivate additional investigations into the functional roles of high frequency neural oscillations across the basal ganglia-thalamocortical motor circuit, similarities and differences across nodal points, and their relationship to motor control in normal and diseased states. These findings also provide rationale for further exploration into how these signals can inform DBS device programming and the development of electrophysiological biomarker-based adaptive stimulation approaches in patients receiving GPi DBS.

## Acknowledgements

We would like to acknowledge the following people for their contributions to this study: Greg Molnar for his help with acquiring equipment components and for helpful insights regarding sensing with directional DBS leads; Ethan Marshall, Sinta Fergus, Stephanie Alberico and Kevin O’Neill for help during data collections; Matthew Dodd and Danielle Koenig for their help collecting clinical rating scores; Tara Palnitkar and Henry Braun for aiding with data collection and image processing, Sommer Huffmaster and Colum MacKinnon for suggestions regarding patient data collection; Jianyu Zhang for assistance with animal preparations; the entire UMN Udall team for critiques and comments on the manuscript.

## References

Aman JE, Johnson LA, Sanabria DE, Wang J, Patriat R, Hill M, et al. Directional deep brain stimulation leads reveal spatially distinct oscillatory activity in the globus pallidus internus of Parkinson’s disease patients. Neurobiol Dis 2020; 139: 104819.

AuYong N, Malekmohammadi M, Ricks-Oddie J, Pouratian N. Movement-Modulation of Local Power and Phase Amplitude Coupling in Bilateral Globus Pallidus Interna in Parkinson Disease [Internet]. Front Hum Neurosci 2018; 12[cited 2019 Mar 11] Available from: https://www.frontiersin.org/articles/10.3389/fnhum.2018.00270/full

Bergman H, Wichmann T, Karmon B, DeLong MR. The primate subthalamic nucleus. II. Neuronal activity in the MPTP model of parkinsonism. J Neurophysiol 1994; 72: 507–520.

Bokil H, Andrews P, Kulkarni JE, Mehta S, Mitra PP. Chronux: A platform for analyzing neural signals. J Neurosci Methods 2010; 192: 146–151.

Boraud T, Bezard E, Guehl D, Bioulac B, Gross C. Effects of L-DOPA on neuronal activity of the globus pallidus externalis (GPe) and globus pallidus internalis (GPi) in the MPTP-treated monkey. Brain Res 1998; 787: 157–160.

Brown P. Oscillatory nature of human basal ganglia activity: relationship to the pathophysiology of Parkinson’s disease. Mov Disord Off J Mov Disord Soc 2003; 18: 357–363.

Burns RS, Chiueh CC, Markey SP, Ebert MH, Jacobowitz DM, Kopin IJ. A primate model of parkinsonism: selective destruction of dopaminergic neurons in the pars compacta of the substantia nigra by N-methyl-4-phenyl-1,2,3,6-tetrahydropyridine. Proc Natl Acad Sci U S A 1983; 80: 4546–4550.

Buzsaki G, Horvath Z, Urioste R, Hetke J, Wise K. High-frequency network oscillation in the hippocampus. Science 1992; 256: 1025–1027.

Du G, Zhuang P, Hallett M, Zhang Y-Q, Li J-Y, Li Y-J. Properties of oscillatory neuronal activity in the basal ganglia and thalamus in patients with Parkinson’s disease. Transl Neurodegener 2018; 7: 17.

Duchin Y, Shamir RR, Patriat R, Kim J, Vitek JL, Sapiro G, et al. Patient-specific anatomical model for deep brain stimulation based on 7 Tesla MRI [Internet]. PLoS ONE 2018; 13[cited 2020 Jun 5] Available from: https://www.ncbi.nlm.nih.gov/pmc/articles/PMC6104927/

Elder CM, Hashimoto T, Zhang J, Vitek JL. Chronic implantation of deep brain stimulation leads in animal models of neurological disorders. J Neurosci Methods 2005; 142: 11–16.

Engel AK, Fries P. Beta-band oscillations—signalling the status quo? Curr Opin Neurobiol 2010; 20: 156–165.

Escobar D, Johnson LA, Nebeck SD, Zhang J, Johnson MD, Baker KB, et al. Parkinsonism and Vigilance: Alteration in neural oscillatory activity and phase-amplitude coupling in the basal ganglia and motor cortex. J Neurophysiol 2017: jn.00388.2017.

Escobar Sanabria D, Johnson LA, Nebeck SD, Zhang J, Johnson MD, Baker KB, et al. Parkinsonism and vigilance: alteration in neural oscillatory activity and phase-amplitude coupling in the basal ganglia and motor cortex. J Neurophysiol 2017; 118: 2654–2669.

Filion M, Tremblay L. Abnormal spontaneous activity of globus pallidus neurons in monkeys with MPTP-induced parkinsonism. Brain Res 1991; 547: 142–151.

Foffani G, Priori A, Egidi M, Rampini P, Tamma F, Caputo E, et al. 300-Hz subthalamic oscillations in Parkinson’s disease. Brain 2003; 126: 2153–2163.

Froemke RC, Dan Y. Spike-timing-dependent synaptic modification induced by natural spike trains. Nature 2002; 416: 433–438.

Hammond C, Bergman H, Brown P. Pathological synchronization in Parkinson’s disease: networks, models and treatments. Trends Neurosci 2007; 30: 357–364.

Hashimoto T, Elder CM, Okun MS, Patrick SK, Vitek JL. Stimulation of the subthalamic nucleus changes the firing pattern of pallidal neurons. J Neurosci Off J Soc Neurosci 2003; 23: 1916–1923.

Hellerbach A, Dembek TA, Hoevels M, Holz JA, Gierich A, Luyken K, et al. Directional Orientation Detection of Segmented Deep Brain Stimulation Leads: A Sequential Algorithm Based on CT Imaging. Stereotact Funct Neurosurg 2018; 96: 335–341.

Hirschmann J, Butz M, Hartmann CJ, Hoogenboom N, Özkurt TE, Vesper J, et al. Parkinsonian Rest Tremor Is Associated With Modulations of Subthalamic High-Frequency Oscillations. Mov Disord 2016; 31: 1551–1559.

Horn A, Neumann W-J, Degen K, Schneider G-H, Kühn AA. Toward an electrophysiological “sweet spot” for deep brain stimulation in the subthalamic nucleus. Hum Brain Mapp 2017; 38: 3377–3390.

Jun JJ, Steinmetz NA, Siegle JH, Denman DJ, Bauza M, Barbarits B, et al. Fully integrated silicon probes for high-density recording of neural activity. Nature 2017; 551: 232–236.

Koop MM, Andrzejewski A, Hill BC, Heit G, Bronte-Stewart HM. Improvement in a quantitative measure of bradykinesia after microelectrode recording in patients with Parkinson’s disease during deep brain stimulation surgery. Mov Disord 2006; 21: 673–678.

Kühn AA, Kupsch A, Schneider G-H, Brown P. Reduction in subthalamic 8-35 Hz oscillatory activity correlates with clinical improvement in Parkinson’s disease. Eur J Neurosci 2006; 23: 1956–1960.

Kühn AA, Tsui A, Aziz T, Ray N, Brücke C, Kupsch A, et al. Pathological synchronisation in the subthalamic nucleus of patients with Parkinson’s disease relates to both bradykinesia and rigidity. Exp Neurol 2009; 215: 380–387.

Levy R, Dostrovsky JO, Lang AE, Sime E, Hutchison WD, Lozano AM. Effects of apomorphine on subthalamic nucleus and globus pallidus internus neurons in patients with Parkinson’s disease. J Neurophysiol 2001; 86: 249–260.

Lofredi R, Neumann W, Brücke C, Huebl J, Krauss JK, Schneider G, et al. Pallidal beta bursts in Parkinson’s disease and dystonia. Mov Disord 2019; 34: 420–424.

López-Azcárate J, Tainta M, Rodríguez-Oroz MC, Valencia M, González R, Guridi J, et al. Coupling between Beta and High-Frequency Activity in the Human Subthalamic Nucleus May Be a Pathophysiological Mechanism in Parkinson’s Disease. J Neurosci 2010; 30: 6667–6677.

Meidahl AC, Tinkhauser G, Herz DM, Cagnan H, Debarros J, Brown P. Adaptive Deep Brain Stimulation for Movement Disorders: The Long Road to Clinical Therapy. Mov Disord 2017; 32: 810–819.

Özkurt TE, Butz M, Homburger M, Elben S, Vesper J, Wojtecki L, et al. High frequency oscillations in the subthalamic nucleus: a neurophysiological marker of the motor state in Parkinson’s disease. Exp Neurol 2011; 229: 324–331.

Patriat R, Cooper SE, Duchin Y, Niederer J, Lenglet C, Aman J, et al. Individualized tractography-based parcellation of the globus pallidus pars interna using 7T MRI in movement disorder patients prior to DBS surgery. NeuroImage 2018; 178: 198–209.

Petersson P, Halje P, Cenci MA. Significance and Translational Value of High-Frequency Cortico-Basal Ganglia Oscillations in Parkinson’s Disease. J Park Dis 2019; 9: 183–196.

Priori A, Foffani G, Pesenti A, Bianchi A, Chiesa V, Baselli G, et al. Movement-related modulation of neural activity in human basal ganglia and its L-DOPA dependency: recordings from deep brain stimulation electrodes in patients with Parkinson’s disease. Neurol Sci 2002; 23: s101–s102.

Ray NJ, Jenkinson N, Wang S, Holland P, Brittain JS, Joint C, et al. Local field potential beta activity in the subthalamic nucleus of patients with Parkinson’s disease is associated with improvements in bradykinesia after dopamine and deep brain stimulation. Exp Neurol 2008; 213: 108–113.

Shreve LA, Velisar A, Malekmohammadi M, Koop MM, Trager M, Quinn EJ, et al. Subthalamic oscillations and phase amplitude coupling are greater in the more affected hemisphere in Parkinson’s disease. Clin Neurophysiol 2017; 128: 128–137.

Silberstein P, Kühn AA, Kupsch A, Trottenberg T, Krauss JK, Wöhrle JC, et al. Patterning of globus pallidus local field potentials differs between Parkinson’s disease and dystonia. Brain 2003; 126: 2597–2608.

Tass PA. A model of desynchronizing deep brain stimulation with a demand-controlled coordinated reset of neural subpopulations. Biol Cybern 2003; 89: 81–88.

Tinkhauser G, Pogosyan A, Debove I, Nowacki A, Shah SA, Seidel K, et al. Directional Local Field Potentials: A Tool to Optimize Deep Brain Stimulation. Mov Disord 2018; 33: 159–164.

Tsiokos C, Hu X, Pouratian N. 200–300Hz movement modulated oscillations in the internal globus pallidus of patients with Parkinson’s Disease. Neurobiol Dis 2013; 54: 464–474.

Tsiokos C, Malekmohammadi M, AuYong N, Pouratian N. Pallidal low β-low γ phase-amplitude coupling inversely correlates with Parkinson disease symptoms. Clin Neurophysiol 2017; 128: 2165–2178.

Vitek JL, Bakay RA, Hashimoto T, Kaneoke Y, Mewes K, Zhang JY, et al. Microelectrode-guided pallidotomy: technical approach and its application in medically intractable Parkinson’s disease. J Neurosurg 1998; 88: 1027–1043.

Vitek JL, Jain R, Chen L, Tröster AI, Schrock LE, House PA, et al. Subthalamic nucleus deep brain stimulation with a multiple independent constant current-controlled device in Parkinson’s disease (INTREPID): a multicentre, double-blind, randomised, sham-controlled study. Lancet Neurol 2020; 19: 491–501.

Vitek JL, Johnson LA. Understanding Parkinson’s disease and deep brain stimulation: Role of monkey models. Proc Natl Acad Sci U S A 2019; 116: 26259–26265.

